# Differential acclimation kinetics of the two forms of Type IV chromatic acclimaters occurring in marine *Synechococcus* cyanobacteria

**DOI:** 10.1101/2023.07.11.548510

**Authors:** Louison Dufour, Bastian Gouriou, Julia Clairet, Morgane Ratin, Laurence Garczarek, Frédéric Partensky

**Affiliations:** Sorbonne Université, CNRS, UMR 7144 Adaptation and Diversity in the Marine Environment (AD2M), Station Biologique de Roscoff (SBR), Roscoff, France

**Author notes:** Correspondence: Corresponding Author: F. Partensky.

**Keywords:** Marine picocyanobacteria, *Synechococcus*, chromatic acclimation

## Abstract

*Synechococcus* is one of two most abundant phytoplanktonic organisms of the Ocean and also displays the widest variety of pigmentation of all marine oxyphotrophs, which makes it ideally suited to colonize the variety of spectral niches occurring in the upper-lit layer of oceans. Seven *Synechococcus* pigment types (PTs) have been described based on the composition and chromophorylation of their light-harvesting complexes, called phycobilisomes (PBS). The most sophisticated and abundant *Synechococcus* PT (3d) gathers cells capable of Type IV chromatic acclimation (CA4), i.e. to reversibly modify the ratio of the blue light-absorbing phycourobilin (PUB) to the green light-absorbing phycoerythrobilin (PEB) in PBS in order to match the ambient light color. Although two genetically distinct types of CA4-capable strains, so-called PTs 3dA and 3dB, have been evidenced and found to be equally abundant in the Ocean, reasons for their prevalence in natural *Synechococcus* populations remain obscure. Here, acclimation experiments in different blue to green ratios of representatives of these two PTs showed that in mixed blue-green light conditions, PT 3dB strains displayed significantly higher PUB:PEB ratios than their PT 3dA counterparts. Thus, PTs 3dA and 3dB seem to differ in the ratio of blue to green light required to trigger the CA4 process. Furthermore, shift experiments between 100% BL and 100% GL conditions, and conversely, also revealed discrepancies in the acclimation pace between the two types of chromatic acclimaters, which may explain their co-occurrence in some blue green light niches.

## INTRODUCTION

Phytoplanktonic cells have an obligate requirement for light to perform photosynthesis. In the marine environment this energy source is highly variable quantitatively and qualitatively not only with depth but also along coast-offshore gradients, leaving aside daily oscillations (Kirk, 1994; Holtrop et al., 2021). This variability has triggered an extensive structural and pigment diversification of phytoplankton light-harvesting antennae. These pigment-protein complexes enable cells to considerably enhance the amount and wavelength range of photosynthetically active radiations reaching photosystems. Despite their apparent simplicity compared to algae and higher plants, cyanobacteria possess the most sophisticated form of antennae known in oxyphototrophs, the phycobilisomes (PBS). PBS are huge water-soluble complexes composed of six to eight rods radiating around a central core. Both core and rods are constituted of phycobiliproteins that bind highly fluorescent open-chain tetrapyrroles, called phycobilins (Sidler, 1994). While the PBS core is always made of allophycocyanin and is highly conserved, PBS rods display a very large structural flexibility, since they can be made of phycocyanin (PC) only, or of PC and one or two phycoerythrin (PE) types, PE-I and PE-II (Ong and Glazer, 1991; Six et al., 2007). Additionally, each phycobiliprotein type can bind up to three different kinds of chromophores (or phycobilins). The ultimate degree of sophistication is the capacity for some cyanobacterial cells to modify the composition of their PBS in response to changes in the ambient light color. This process called chromatic acclimation (CA) — the initial term was actually ‘chromatic adaptation’ but was recently replaced in order to best suit this physiological process (Shukla et al., 2012) — was first observed in the early 20^th^ century in freshwater cyanobacteria shifted from red to green light (Engelmann, 1902; Gaidukov, 1903). It was later on attributed to changes in the phycobiliprotein composition of PBS rods: in red light, rods are entirely composed of PC and cells look green, whereas in green light PC is restricted to the base of the rods and the distal part is made of PE, causing cells to exhibit a bright red color (Boresch, 1922). This type of complementary chromatic acclimation (also called Type 3 chromatic acclimation or CA3) is one among the six different types of CA known so far in Cyanobacteria (Tandeau de Marsac, 1977; Sanfilippo et al., 2019a). While CA2 —a simple type of CA where PE production is induced in GL, generating longer PBS rods, and repressed in red light— and CA3 occur mainly in freshwater and brackish cyanobacteria, CA4 is the main type occurring in the open ocean and is specific of marine *Synechococcus* cyanobacteria (Palenik, 2001; Everroad et al., 2006; Humily et al., 2013). Contrary to CA2 and CA3, CA4 does not involve changes in the phycobiliprotein composition of PBS rods, but in their chromophore composition. Indeed, in response to shifts between green light (GL) and blue light (BL), CA4-capable cells can modify the relative amount of the two chromophores bound to PE-I and PE-II, in order to match the predominant ambient light color. In BL they exhibit a high ratio of the BL-absorbing phycourobilin (PUB, *A*_max_ = 495 nm) to the GL-absorbing phycoerythrobilin (PEB, *A*_max_ = 550 nm), and conversely in GL (Everroad et al., 2006; Shukla et al., 2012). Variations in PUB:PEB ratio are generally assessed by measuring the relative ratio of whole cell fluorescence excitation at 495 and 545 nm (Exc_495:545_) with emission at 580 nm, which in CA4-capable strains changes from 0.6-0.7 in GL to 1.6-1.7 in BL (Palenik, 2001; Everroad et al., 2006). In the nomenclature of *Synechococcus* pigment types (PTs) established by (Six et al., 2007), and later modified by (Humily et al., 2013), chromatic acclimaters are classified as ‘PT 3d’ cells, meaning that they possess PBS rods made of PC, PE-I and PE-II (a feature shared by all PT 3 strains) and can modify their PUB:PEB ratio. In contrast PTs 3a, 3b and 3c have a constitutively low, medium and high PUB:PEB ratio, respectively. Additionally, two genetically different types of CA4-capable strains have been described: PTs 3dA and 3dB (Humily et al., 2013). They possess a small genomic island involved in the CA4 process (CA4-A and CA4-B), both forms differing genetically and structurally (Humily et al., 2013; Sanfilippo et al., 2019b; Grébert et al., 2021). Although long overlooked, CA4 appears to be an ecologically important process for marine *Synechococcus*, since CA4-capable cells were shown to account for more than 40% of the whole *Synechococcus* population along the *Tara* Oceans expedition transect (Grébert et al., 2018). Moreover, PTs 3dA and 3dB were equally abundant (22.6% and 18.9%, respectively) but distributed in complementary light niches in the field, the former predominating in temperate and high latitude waters, while the latter were more abundant in warm waters.

The emergence and maintenance of two CA4 types over the course of evolution as well as their differential distribution in the environment suggests that both CA4 types may not be as phenotypically equivalent as previously suggested (Humily et al., 2013). To check this hypothesis, we acclimated three representatives of PTs 3dA and 3dB in batch culture under two conditions of temperature, two irradiances and five light colors to compare their growth rates and PBS characteristics. Furthermore, we performed shifts at different light intensities from BL to GL (and conversely) to compare CA4 kinetics between five strains of each PT 3d. Our study indicates that PT 3dA and 3dB strains differ in the BL to GL ratios necessary to trigger the CA4 process. Moreover, we revealed discrepancies in the acclimation pace between the two types of chromatic acclimaters.

## MATERIALS AND METHODS

### Biological material and culture conditions

Ten *Synechococcus* strains, of which five PT 3dA and five PT 3dB representative strains (Table 1), were retrieved from the Roscoff Culture Collection (https://roscoff-culture-collection.org/). These strains were isolated from very diverse environments and were selected based on genome availability (Doré et al., 2020) and clade affiliation in order to have representatives from all five major clades (I to IV and CRD1) in the global ocean (Farrant et al., 2016), as well as the CA4-A model strain RS9916 (Shukla et al., 2012; Sanfilippo et al., 2016, 2019b).

**TABLE 1.** Characteristics of the *Synechococcus* strains used in this study.

| Strain name | RCC #<sup>1</sup> | Subcluster<sup>2</sup> | Clade<sup>2</sup> | Subclade<sup>3</sup> | Pigment type<sup>4</sup> | Isolation region |
| --- | --- | --- | --- | --- | --- | --- |
| BIOS-U3-1 | 2533 | 5.1 | CRD1 | n.a. | 3dA | Chile upwelling |
| BL107 | 515 | 5.1 | IV | IVa | 3dA | Balearic Sea |
| MITS9220 | 2571 | 5.1 | CRD1 | n.a. | 3dA | Equatorial Pacific |
| RS9916 | 555 | 5.1 | IX | n.a. | 3dA | Gulf of Aqaba |
| WH8020 | 751 | 5.1 | I | Ia | 3dA | Sargasso Sea |
| A15-62 | 2374 | 5.1 | II | IIa | 3dB | Offshore Mauritania |
| A18-40 | n.a. | 5.1 | III | IIIa | 3dB | Atlantic Ocean |
| MINOS11 | 2319 | 5.3 | n.a. | n.a. | 3dB | Mediterranean Sea |
| PROS-U-1 | 2369 | 5.1 | II | IIh | 3dB | Moroccan upwelling |
| RS9915 | 2553 | 5.1 | III | IIIA | 3dB | Gulf of Aqaba |

<sup>1</sup>Roscoff Culture Collection, <sup>2</sup>Farrant et al. (2016), <sup>3</sup>Mazard et al. (2012), <sup>4</sup>Humily et al. (2013).

Cells were grown in 50 mL flasks (Sarstedt) in PCR-S11 medium (Rippka et al., 2000) supplemented with 1 mM sodium nitrate and pre-acclimated for at least three weeks in continuous light, provided by blue and/or green LEDs (Alpheus) in temperature-controlled chambers.

### Acclimation experiments

A selection of six out of the ten abovementioned strains (BL107, RS9916 and WH8020 for PT 3dA and A15-62, PROS-U-1 and RS9915 for PT 3dB; Table 1) were grown in the following conditions: (i) two temperatures: 18 and 25°C; (ii) two irradiances: low light (LL, 15 μmol photons m^−2^ s^−1^, hereafter µE) and high light (HL, 75 µE); (iii) five light qualities: blue light (100% BL), green light (100% GL), as well as three mixtures of blue to green light: 25% BL – 75% GL, 50% BL – 50% GL and 75% BL – 25% GL. The LEDs spectra were measured in these different conditions using a PG200N Spectral PAR Meter (UPRtek). Each strain was grown in triplicates and inoculated at an initial cell density of 3 × 10^6^ cells mL^−1^. Samples were harvested every day to measure cell concentration and fluorescence parameters by flow cytometry, and once during the exponential phase to measure phycobilin content by spectrofluorimetry (see below).

### Shift experiments

Shift experiments between 100% low BL (LBL) and 100% low GL (LGL), and conversely, as well as the equivalent experiments between 100% high BL (HBL) and 100% high GL (HGL), were performed on all ten *Synechococcus* strains (Table 1), but only at 25°C. Each strain was diluted with fresh medium before the beginning of the experiments and regularly transferred in order to avoid limitation by nutrients. Aliquots were collected two to three times a day, depending on light intensity, to measure phycobilin content by spectrofluorimetry (see below).

### Flow cytometry

Culture aliquots were sampled twice a day, fixed using 0.25% (v/v) glutaraldehyde (grade II, Sigma Aldrich) and stored at −80°C until analysis (Marie et al., 1999). Cell density was determined using a Guava easyCyte flow cytometer equipped with a 488 nm laser and the Guavasoft software (Luminex Corporation). Average fluorescence at 583 nm and 695 nm were used as proxies of phycoerythrin (PE) and chlorophyll *a* (Chl *a*) contents per cell, respectively. Forward scatter (FSC) was used as a proxy of cell diameter. Both fluorescence and FSC signals were normalized to that of standard fluorescent 0.95 μm silica beads.

### Spectrofluorimetry

*In vivo* fluorescence spectra were recorded at 240 nm min^-1^ with slits fixed at 10 nm once during the exponential phase using a spectrofluorimeter FL6500 (Perkin-Elmer). Excitation spectra were acquired between 450 and 560 nm (with an emission set at 580 nm, which corresponds to the maximum emission of PE). Emission spectra were recorded between 550 and 750 nm (with an excitation set at 530 nm, which corresponds to the maximum absorption of PUB). Spectra were monitored and analyzed with the Fluorescence software (Perkin-Elmer). The excitation ratio (Exc_495:550_) was considered as a proxy for PUB:PEB ratio, an indicator of the phycoerythrobilin and phycourobilin content of the PBS rods. The fluorescence emission ratios (Em_560:650_) and (Em_650:680_) were respectively used as proxy of phycoerythrin (PE) to phycocyanin (PC) and PC to PBS terminal acceptor (TA) ratios. Those provided information about the electron transfer efficiency within the PBS, the length of PBS rods and the coupling of the PBS to PSII reaction center chlorophylls.

## RESULTS

### Acclimation experiments

#### Growth rate

The growth rates of all six tested strains were globally higher at 25°C (Fig. 1C and D) than at 18°C (Fig. 1A and B). Similarly, most strains grew significantly faster at HL than LL at 25°C, while little difference between the two irradiances was observed at 18°C. Consequently, maximum growth rates (µ) were reached at 25°C in HL, with all strains but WH8020 achieving more than one cell division per day (µ > 0.69 day^-1^; Fig. 1D). While for any given strain, the growth rate varied little between the different light quality conditions, significant differences were observed between PTs 3dA and 3dB at 18°C and LL under four out of five light quality conditions, with PT 3dA strains growing slightly faster than their PT 3dB counterparts (Fig. 1A). This was also the case for the 100% BL condition at HL and 18°C (Fig. 1B), as well as at 50% BL – 50% GL and 100% GL at LL and 25°C (Fig. 1C). No statistically significant differences between the two PTs were observed at 25°C HL.

**Figure 1:**
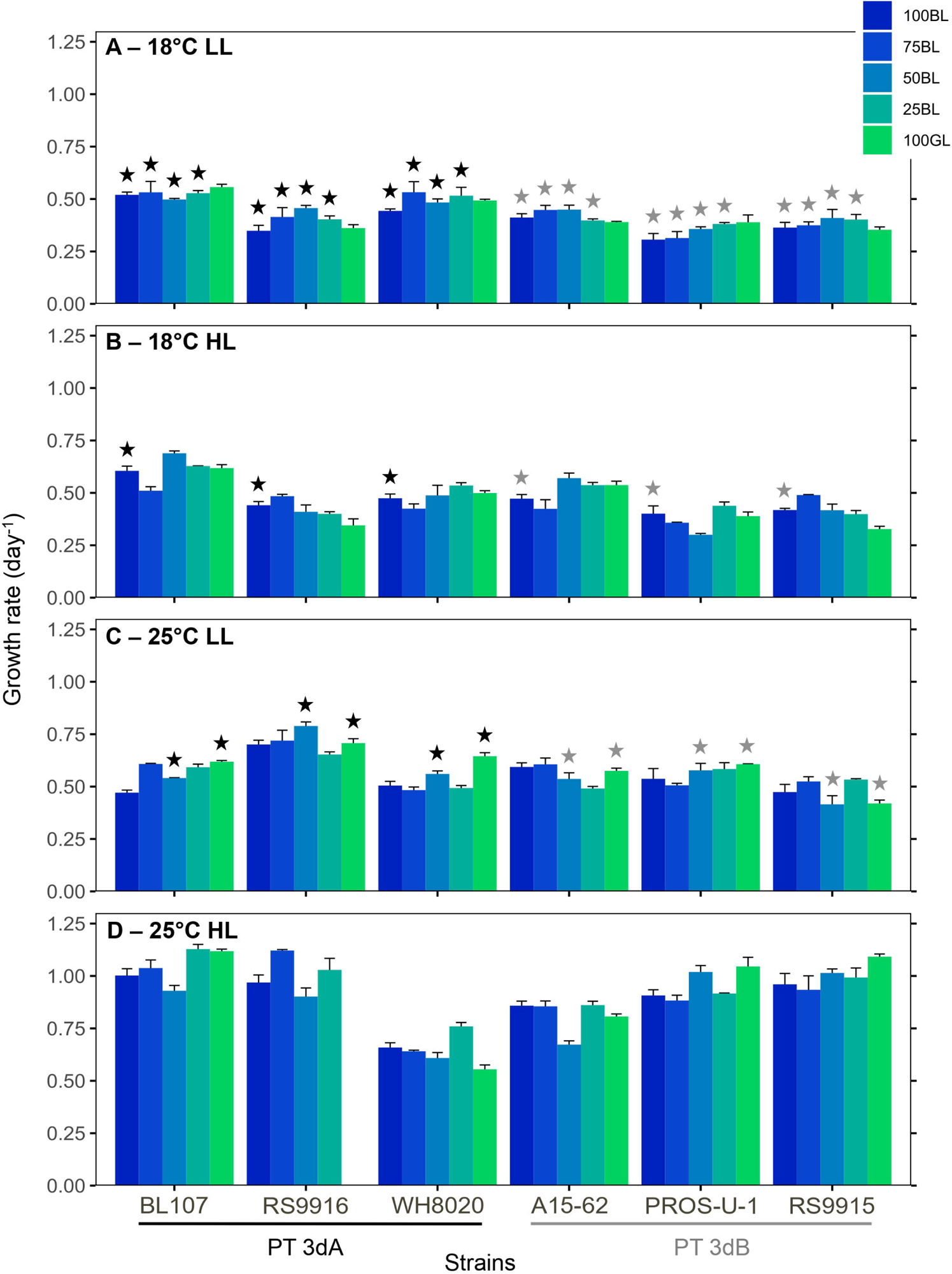
Growth rate of six *Synechococcus* strains grown in different conditions of temperature, light quantity and quality. **(A)** 18°C low light. **(B)** 18°C high light. **(C)** 25°C low light. (**D**) 25°C high light. Histograms represent average and standard deviations of triplicate measurements. Stars above histograms indicate significantly different values between PT 3dA and 3dB represenatives in a given condition of light quality and temperature (Student test or Wilcoxon-Mann-Whitney test, depending on normality and variance homogeneity, p-value < 0.05).

#### Chlorophyll *a* and phycoerythrin fluorescence

A downward trend in flow cytometric red (Chl *a*) and orange (PE) fluorescence signals was observed from 100% BL to 100% GL, regardless of irradiance and temperature (Fig. 2 and 3). Both fluorescence signals were higher in LL (Fig. 2A and C, 3A and C) than in HL (Fig. 2B and D, 3B and D), but also at 25°C (Fig. 2C and D, 3C and D) in comparison to 18°C (Fig. 2A and B, 3A and B). Significant differences between PTs 3dA and 3dB were detected only at LL (Fig. 2A and C, 3A and C). At 18°C, the Chl *a* fluorescence of PT 3dB strains was higher than for PT 3dA cells at 25% BL – 75% GL and 100% GL (Fig. 2A). The same was true for PE fluorescence in all tested light qualities except 100% BL (Fig. 3A). At 25°C, PT 3dB strains exhibited higher Chl *a* fluorescence in the three light mixtures (Fig. 2C) and PE fluorescence in three out of the five light quality conditions (Fig. 3C).

**Figure 2:**
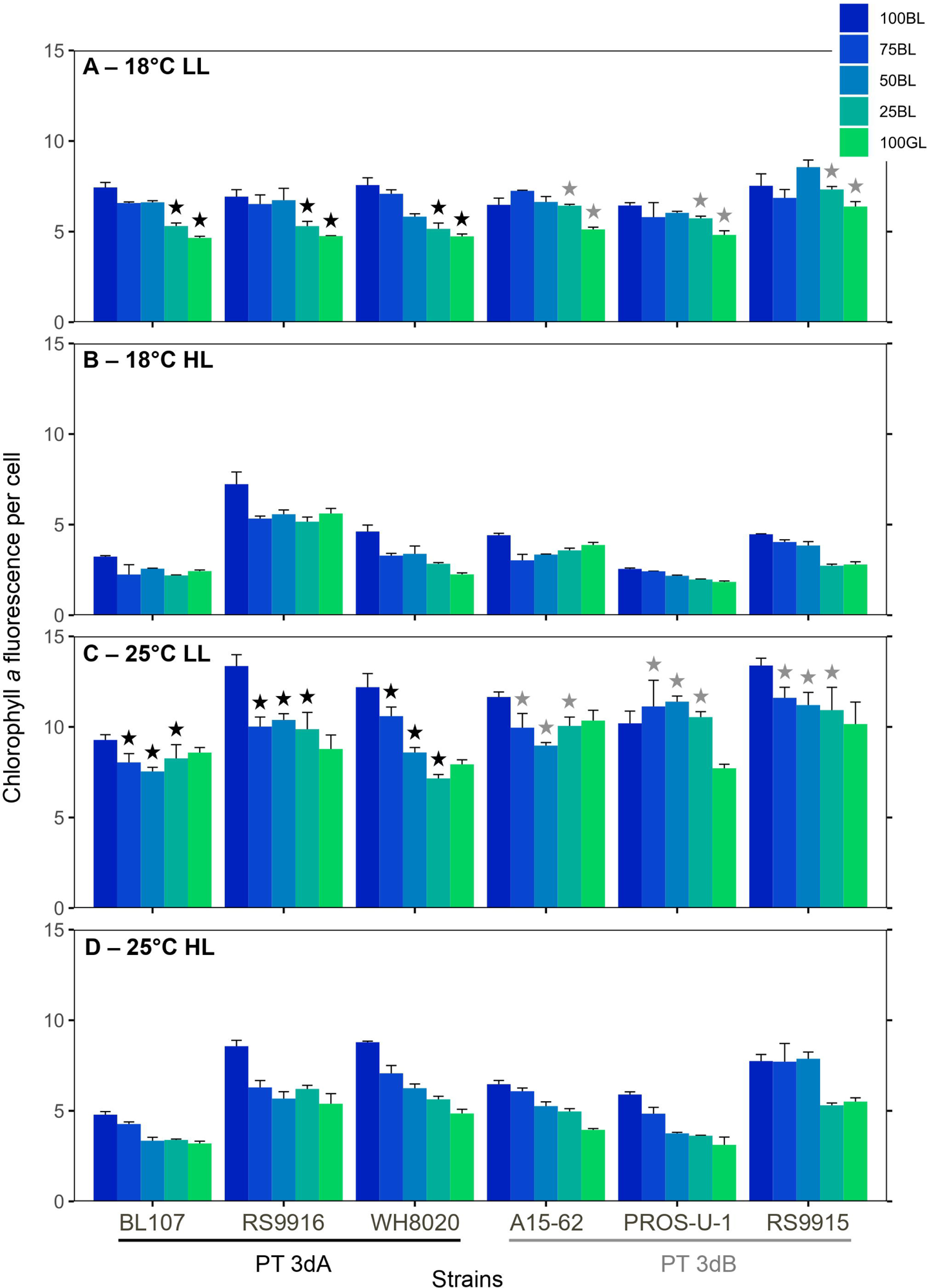
Average flow cytometric chlorophyll *a* fluorescence per cell of six *Synechococcus* strains grown in different conditions of temperature, light quantity and quality. **(A)** 18°C low light. **(B)** 18°C high light. **(C)** 25°C low light. (**D**) 25°C high light. Histograms represent average and standard deviations of triplicate measurements. Stars above histograms indicate significantly different values between PT 3dA and 3dB representatives in a given condition of light quality and temperature (Student test or Wilcoxon-Mann-Whitney test, depending on normality and variance homogeneity, p-value < 0.05).

**Figure 3:**
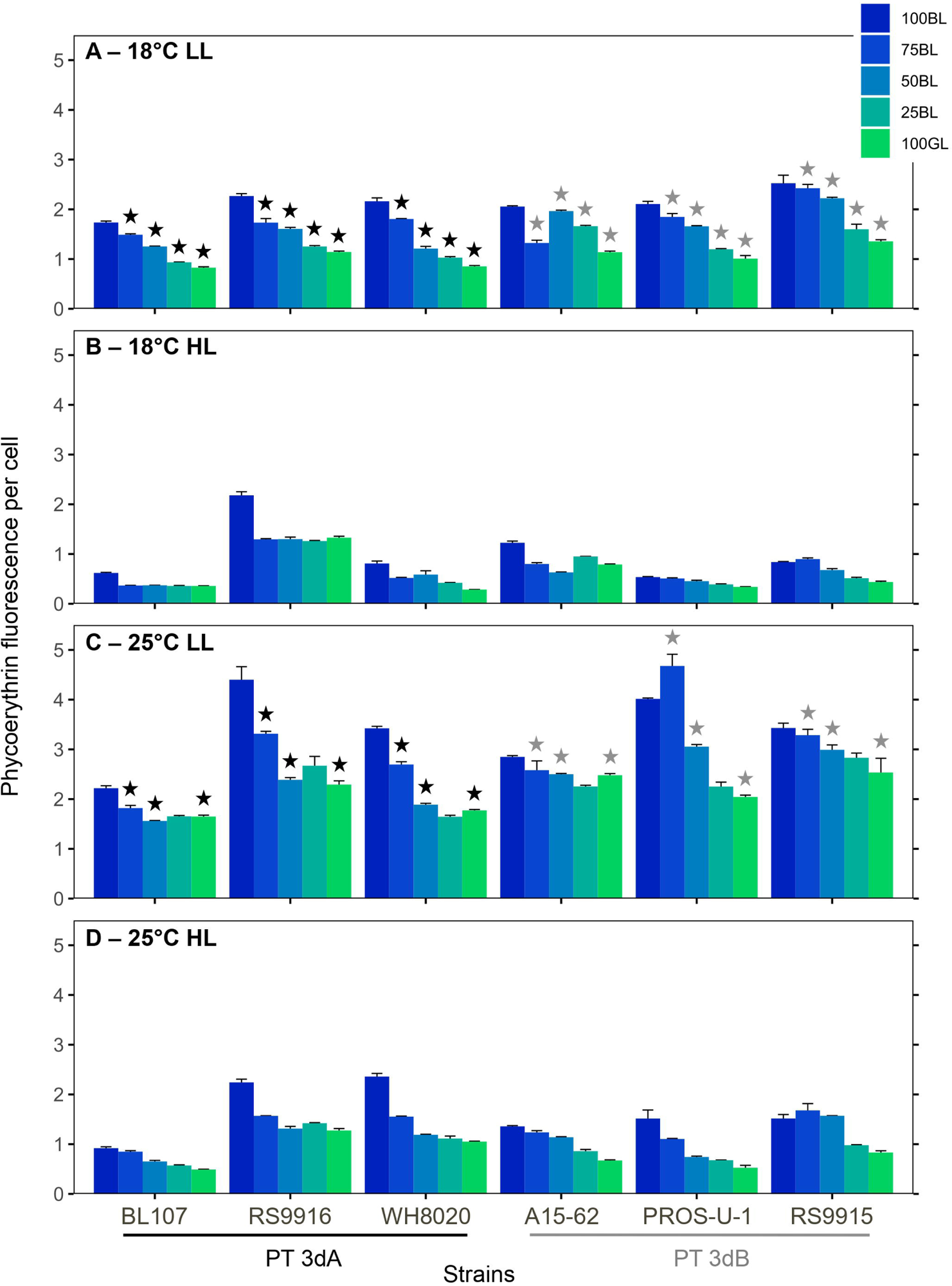
Average flow cytometric phycoerythrin fluorescence per cell of six *Synechococcus* strains grown in different conditions of temperature, light quantity and quality. **(A)** 18°C low light. **(B)** 18°C high light. **(C)** 25°C low light. (**D**) 25°C high light. Histograms represent average and standard deviations of triplicate measurements. Stars above histograms indicate significantly different values between PT 3dA and 3dB representatives in a given condition of light quality and temperature (Student test or Wilcoxon-Mann-Whitney test, depending on normality and variance homogeneity, p-value < 0.05).

#### Phycobilin content

As expected, the Exc_495:550_ fluorescence excitation ratio, a proxy of the PUB:PEB ratio of the cells, decreased from 100% BL to 100% GL in all conditions tested (Fig. 4A to D). With a few exceptions, all strains displayed typical Exc_495:545_ ratios for chromatic acclimaters when acclimated to 100% GL (0.6-0.7) or 100% BL (1.6-1.7; (Humily et al., 2013). It is important to note that the LEDs that were used to get the 100% GL condition actually peaked at 515 nm, which is at the blue edge of the green wavelength range (Supplemental Fig. 1). Yet, the fact that all tested CA4-capable strains exhibited the lowest possible Exc_495:545_ ratio shows that they all ‘senses’ this light quality as being GL.

**Figure 4:**
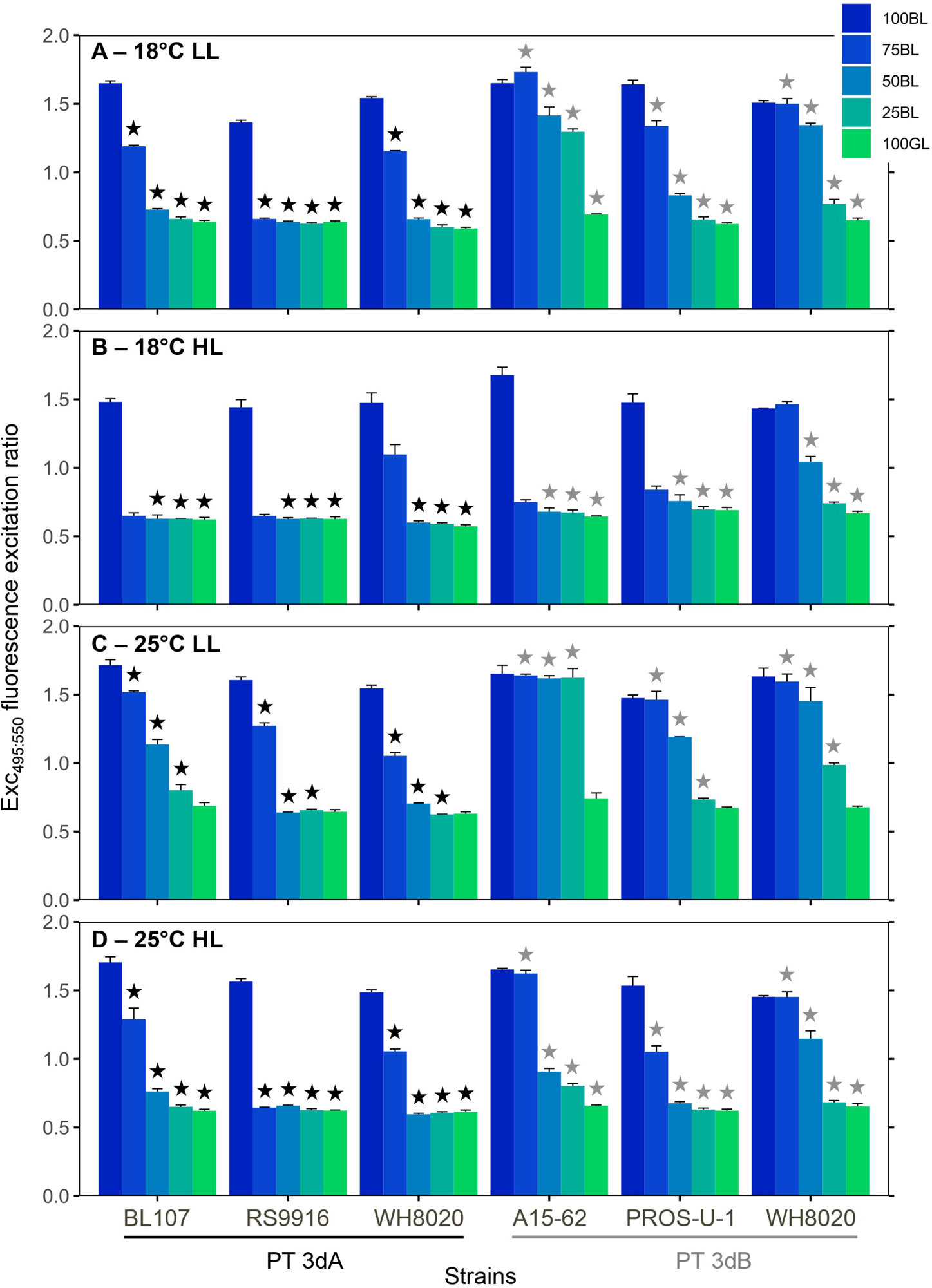
Exc_495:550_ fluorescence excitation ratio (a proxy of PUB:PEB ratio) of six *Synechococcus* strains grown in different conditions of temperature, light quantity and quality. **(A)** 18°C low light. **(B)** 18°C high light. **(C)** 25°C low light. (**D**) 25°C high light. Histograms represent average and standard deviations of triplicate measurements. The stars above histograms indicate significantly different values between PT 3dA and 3dB representatives in a given condition of light quality and temperature (Student test or Wilcoxon-Mann-Whitney test, depending on normality and variance homogeneity, p-value < 0.05).

Statistical tests revealed that PT 3dB representatives almost always displayed significantly higher Exc_495:545_ ratios than their PT 3dA counterparts, except in 100 % BL (Fig. 4A to D and Supplemental Fig. 1). Comparison between LL and HL showed that under LL, most cells exhibited progressively decreasing Exc_495:545_ ratios in the three mixed blue-green light conditions (Fig. 4A and C), whereas in HL there was generally a more abrupt shift to a ratio typical of 100% GL acclimation (Fig. 4B and D). As concerns temperature, its effect on PUB:PEB ratios appeared to be much less marked.

#### Phycobiliprotein content

Two different patterns were identified concerning the variations of Em_560:650_ fluorescence emission ratio, a proxy of PBS PE:PC ratio. Indeed, a decrease in this ratio was most often observed from 100% BL to 100% GL in PT 3dA strains, while it remained fairly stable in PT 3dB representatives, whatever the light color (Fig. 5A to D). The effects of temperature and irradiance were much less clear since the Em_560:650_ ratio did not vary dramatically with these two parameters. Statistically significant differences between PTs 3dA and 3dB were found in a number of conditions, but in contrast to Exc_495:545_ ratios, PT 3dA cells exhibited the highest Em_560:650_ values.

**Figure 5:**
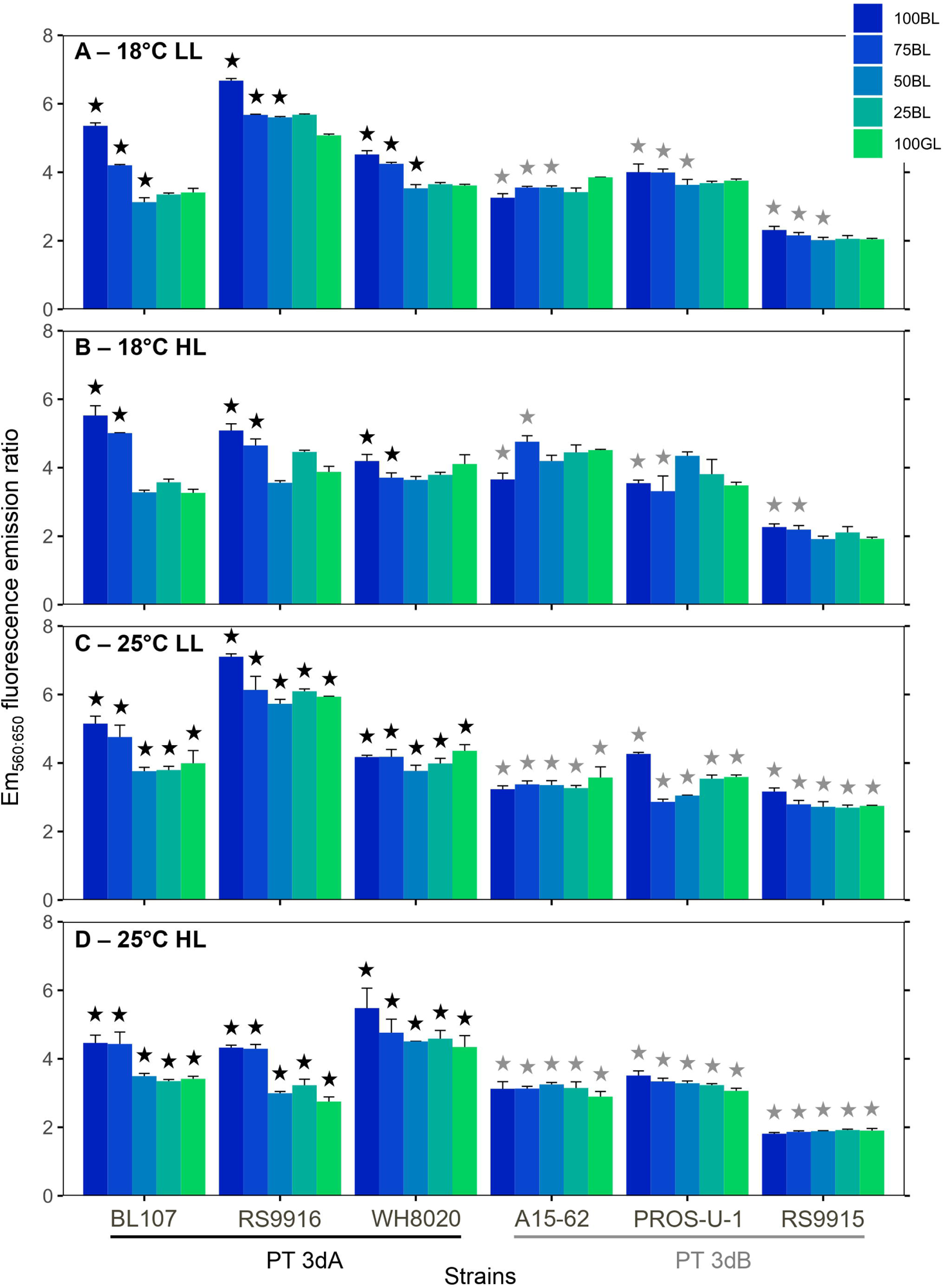
Em_560:650_ fluorescence emission ratios (a proxy of PE to PC ratio) of six *Synechococcus* strains grown in different conditions of temperature, light quantity and quality. **(A)** 18°C low light. **(B)** 18°C high light. **(C)** 25°C low light. (**D**) 25°C high light. Histograms represent average and standard deviations of triplicate measurements. The stars above histograms indicate significantly different values between PT 3dA and 3dB representatives in a given condition of light quality and temperature (Student test or Wilcoxon-Mann-Whitney test, depending on normality and variance homogeneity, p-value < 0.05).

As concerns the Em_650:680_ fluorescence emission ratio, a proxy of the PC:TA ratio, little variations were observed for a given strain, with values always between 1 and 2 whatever the conditions, except for RS9916 that displayed values above 2 in several conditions (Supplemental Fig. 2A to D). Yet, PT 3dA strains had significantly higher values than their PT 3dB counterparts almost exclusively in HL (Supplemental Fig. 2C and D).

#### Cell diameter

In general, the FSC, a proxy of cell diameter, varied little with light quality (Supplemental Fig. 3A to D). As concerns light intensity and temperature, different patterns were observed. For example, the diameter of BL107 seemed to remain constant, regardless of the tested conditions. In others strains, such as PROS-U-1 and RS9915, notable variations in FSC were observed with minimum FSC values achieved for both strains in HL and at 25°C (Supplemental Fig. 3D). Finally, the greatest variations were observed in RS9916 (> 3 times), as well as A15-62 and WH8020 (> 2.5 times), with maximum values in HL but at 18 or 25 °C depending on the strains (Supplemental Fig. 3B and D). Statistical tests showed that PT 3dB cells had significantly larger diameter than PT 3dA strains in some conditions of LL, at 18 and 25°C (Supplemental Fig. 3A and C).

### Shift experiments

Shift experiments from GL to BL, and reverse shifts, in either LL or HL conditions were carried out with five PT 3dA and five PT 3dB representatives (Table 1). As expected from a previous study (Humily et al., 2013), the BIOS-U3-1 strain and another representative of the CRD1 clade (MITS9220) did not fully acclimate to HBL and remained stuck at an Exc_495:545_ ratio of about 1.2 (Supplemental Fig. 4B and D). We therefore compared the kinetics and amplitude of Exc_495:545_ variations between PTs 3dA and 3dB with respect to the initial value without including the CRD1 members (Fig. 6A to D). As expected, PT 3dA and 3dB cells exhibited significantly slower kinetics of chromatic acclimation in LL than in HL since CA4 took about seven days in LL and four days in HL. Furthermore, comparison between the two PTs for a given shift showed that the acclimation kinetics was significantly faster for PT 3dA than PT 3dB from LGL to LBL (Fig. 6A). In contrast, PT 3dB strains exhibited faster acclimation kinetics than their PT 3dA counterparts during the shift from HBL to HGL (Fig. 6D). It is worth noting that, although not significant, similar trends were observed for the two other shift conditions (Fig. 6B and C).

**Figure 6:**
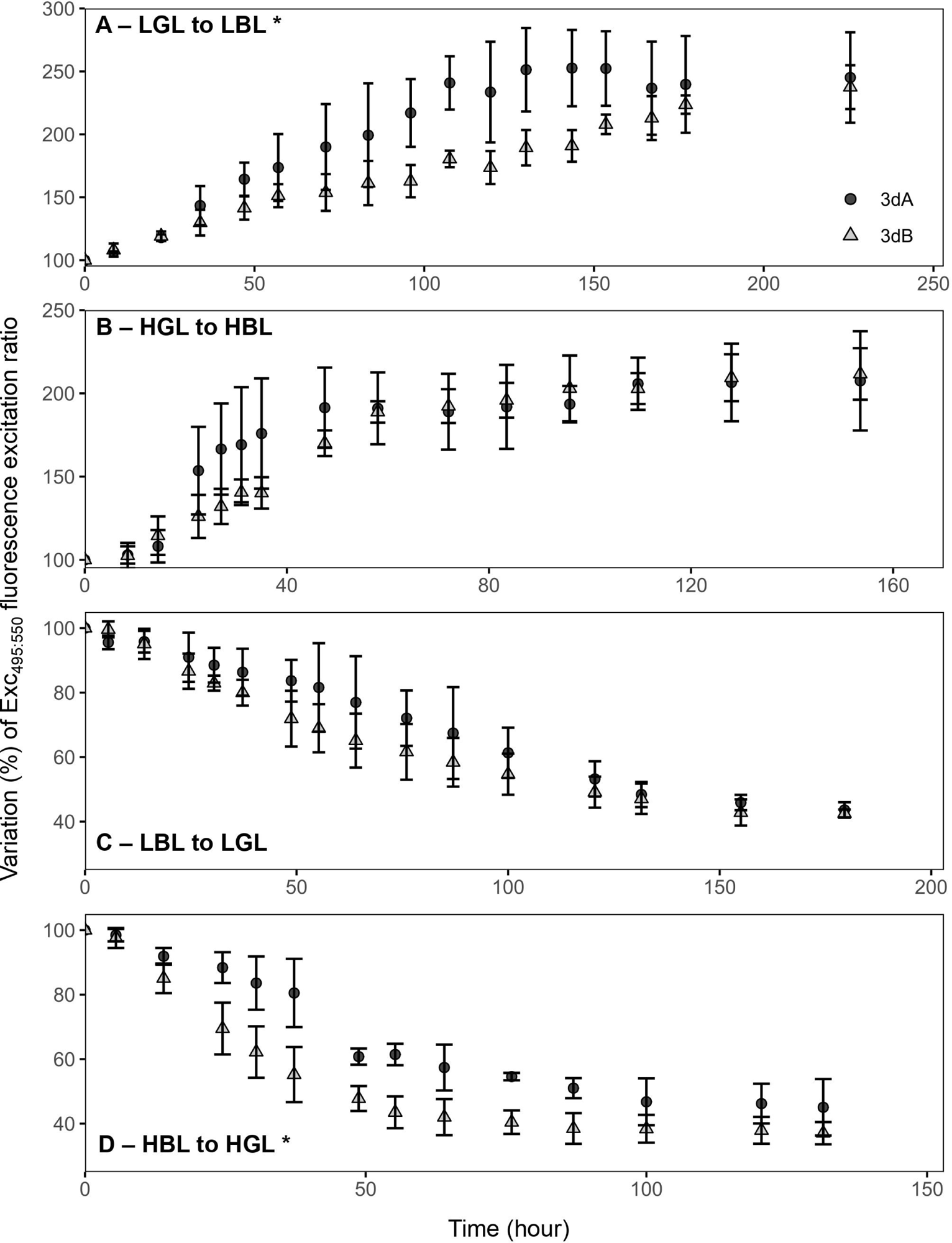
Variation of PUB:PEB compared to initial value of the ten *Synechococcus* strains after an abrupt shift of light quality. **(A)** LGL to LBL. **(B)** HGL to HBL. **(C)** LBL to LGL. (**D**) HBL to HGL. Each point represents average and standard deviations of three to five measurements. The stars indicate significantly different values between PT 3dA and 3dB representatives (Wilcoxon-Mann-Whitney test, p-value < 0.05).

## DISCUSSION

Variations in the underwater light field occur in the upper-lit layer of oceans in response to vertical and horizontal gradients of physico-chemical parameters. These create a number of light niches, to which the different coexisting phytoplankton species are more or less well fitted, depending on their intrinsic pigment content. Recently, (Holtrop et al., 2021) found that the open ocean shelters five different spectral niches, based on the absorption properties of water molecules as well as the variable concentrations of colored dissolved organic matter and non-algal particles. Thanks to their specific antenna complexes binding divinyl derivatives of Chl *a* and *b* (Ralf and Repeta, 1992), *Prochlorococcus* cells appear to be well adapted to the violet niche (401-449 nm), which corresponds to the central oceanic gyres. *Synechococcus* blue light specialists, i.e. cells displaying a high PUB:PEB ratio, are well suited to the blue niche (449-514 nm), which encompasses most other marine areas up to polar waters. As concerns green light specialists, i.e. cells displaying a low PUB:PEB ratio, they preferentially thrive in the green niche (514-605 nm), which is essentially found in ocean border areas (Holtrop et al., 2021). In this context, *Synechococcus* cells capable of CA4, i.e. to match their PUB:PEB ratio to the ambient light color in order to optimize photon collection, expectedly colonize both the blue and green niches where they can constitute a large part of *Synechococcus* population, especially at high latitude (Xia et al., 2017; Grébert et al., 2018). The occurrence of two genetically distinct types of CA4-capable *Synechococcus* cells (Humily et al., 2013; Grébert et al., 2018, 2021, 2022) however made us wonder whether they occupy the very same spectral niche in the marine environment or not. To tackle this question, we looked at the interplay between light quality, light quantity as well as temperature in order to try finding out possible slight differences in pigment phenotypes and/or acclimation kinetics between PTs 3dA and 3dB.

Comparisons of PT 3dA and 3dB cells acclimated to various light qualities ranging from 100% BL to 100% GL at two temperatures and two irradiance levels revealed a number of differences. One of the most striking one was that, in all tested light quality conditions except 100% BL, PT 3dB strains displayed significantly higher Exc_495:545_ ratios than their PT 3dA counterparts, independently of temperature or irradiance levels (Fig. 4). The observed differences are seemingly due to a more progressive decrease of the Exc_495:545_ ratio in intermediate light colors between 100% BL to 100% GL, as well as a slightly higher Exc_495:545_ ratio in the 100% GL condition in PT 3dB than PT 3dA representatives. Indeed, PT 3dA strains were more frequently found in the GL-than the BL-acclimated state with some of them, such as RS9916 in most conditions, even shifting their Exc_495:545_ ratio to the BL-acclimated state only in 100% BL (Fig. 4). In contrast, PT 3dB cells tended to longer remain in the BL-acclimated state, even when the proportion of green light was important, an extreme case being A15-62 at 25°C LL that shifted to the GL-acclimated Exc_495:545_ ratio only in 100% GL (Fig. 4C). Interestingly, it has been hypothesized that the ancestral PT 3dA genotype may have derived from a former GL-specialist having acquired the CA4 capacity by integrating a CA4-A island, while the PT 3dB genotype may have been derived from a former BL specialist having integrated a CA4-B island (Sanfilippo et al., 2019a; Grébert et al., 2021). Our results are consistent with this hypothesis as they suggest that PT 3dA strains need a large proportion of blue photons to induce the CA4-A response, while on the contrary PT 3dB cells require a large proportion of green photons to induce the CA4-B response. In other words, the two PTs seemingly differ in the BL:GL ratio necessary to trigger the CA4 process.

The molecular basis of the difference in PUB:PEB ratio between CA4-capable cells fully acclimated to either BL or GL has been well documented, and was found to be the same in both the PT 3dA model strain RS9916 (Shukla et al., 2012; Sanfilippo et al., 2016, 2019b) and the PT 3dB model strain A15-62 (Grébert et al., 2021). Both CA4-A and -B processes indeed consist in an exchange of one out of the five phycobilins bound to the α-PE-II subunit (Cys-139) and two out of the six phycobilins bound to the α-PE-II subunit (Cys-83 and Cys-140). More precisely, PUB molecules are bound to these positions in BL and PEB in GL. However, the observation of intermediate Exc_495:545_ ratios in mixed blue-green light conditions (Fig. 4; see also (Sanfilippo et al., 2019b) remains difficult to interpret, as it may translate different, but not mutually exclusive, sources of variability. Indeed, this observation may be explained by (i) heterogeneous populations of *Synechococcus* cells with PBS either fully acclimated to BL or to GL; (ii) homogeneous populations of *Synechococcus* cells all having different phycobilins at the three swing sites; (iii) individual cells containing PBS with different chromophorylation states; and/or (iv) heterogeneity in phycobiliprotein chromophorylation within single PBS, i.e. PBS having rods with different chromophorylation states. Unfortunately, there is currently no simple experimental way to demonstrate which of these hypotheses is most likely.

In contrast to light quality, both temperature and irradiance seemingly had little effects on the Exc_495:545_ ratios. In the HL condition, growth rates of all six strains used for acclimation experiments were significantly higher at 25°C (the optimal growth temperature of most marine *Synechococcus* strains tested so far), than at 18°C (Fig. 1), consistent with previous literature (Pittera et al., 2014; Breton et al., 2020; Doré et al., 2020; Ferrieux et al., 2022). Yet, the positive effect of increased temperature on growth rate was much less important at LL than HL, whatever the strain, showing that temperature and irradiance have a synergistic effect on growth of CA4-capable strains, as previously observed in the model *Synechococcus* PT 3a strain WH7803 (Guyet et al., 2020). Of note, whatever the temperature, all CA4-capable strains were also able of classical photoacclimation, i.e. to adjust the surface of photosynthetic membranes to reduce the incoming photon flux (Kana and Glibert, 1987; Moore et al., 1995; Six et al., 2004), as shown by the strong decrease of both Chl *a* and PE contents per cell between LL and HL (Fig. 2 and 3). Interestingly, both parameters also slightly decreased in most strains in response to progressive changes in light quality from 100% BL to 100% GL, suggesting that cells used GL more efficiently than BL. They thus needed to slightly adjust the surface of photosynthetic membranes (i.e. both photosystems and PBS) in BL in order to maximize the collection of available photons and maintain their growth rate, which in contrast to pigment ratios did not significantly vary with light quality (Fig. 1). Of note, a direct comparison of the excitation spectra of CA4-capable strains with LEDs spectra in the different light quality conditions (Supplemental Fig. 1) showed that the light provided by blue LEDs, which peaked at 475 nm, excited both Chl *a* (*A*_max_ = 440 nm) and PUB (*A*_max_ = 495 nm), but not PEB (*A*_max_ = 550 nm). In GL, LEDs emission at 515 nm excited both PUB and PEB (Supplemental Fig. 1). In this context, (Lovindeer et al., 2021) also recently showed that CA4-capable *Synechococcus* strains were characterized by much higher light use efficiency in GL than in BL, often associated with faster growth rate. The latter difference with our results can be explained by the fact that the blue LEDs used in this previous study had a peak at 440 nm, mainly absorbed by Chl *a* (*A*_max_ = 440 nm) but very poorly by PUB (*A*_max_ = 495 nm), resulting in strong limitation of the production of chemical energy by photosynthesis in their BL condition.

Another parameter that differentiated PTs 3dA and 3dB is the Em_560:650_ fluorescence emission ratio, which is often interpreted as a proxy for the PE:PC ratio. This was significantly higher in the PT 3dA strains, especially in the bluest light conditions (Fig. 5). Indeed, this ratio tended to decrease from BL to GL in PT 3dA strains, while it was more stable in PT 3dB representatives. Since PE fluorescence per cell decreased in PT 3dA from 100% BL to 100% GL (Fig. 3) and the PC:TA ratio was somewhat constant (Supplemental Fig. 2), this may indicate a lower within-rod energy transfer efficiency than in their PT 3dB counterparts. Another alternative is that PBS rods were progressively shortened from 100% BL to 100% GL in PT 3dA cells but not in PT 3dB. Indeed, the distal PUB-rich PE-II hexamers were previously shown to be eliminated as a result of photoacclimation in *Synechococcus* sp. WH8102 (Six et al., 2004). In PT 3dB representatives, the PE:PC ratio stability, associated with decreasing PE fluorescence, rather suggests that other kinds of structural changes occurred, such as a reduction in the PBS number per cell or in thylakoidal surface area.

Finally, shifts experiment in both LL and HL conditions between 100% BL and 100% GL conditions, and conversely, confirmed previous observations of (Humily et al., 2013) that PT 3dA cells generally exhibit more variable acclimation kinetics than PT 3dB, especially in HL conditions. Indeed, two PT 3dA strains belonging to the CRD1 clade (BIOS-U3-1 and MITS9220) were stuck at a Exc_495:550_ ratio of around 1.2 in HBL (corresponding to ‘phenotypic group 2’ in (Humily et al., 2013). Moreover, MITS9220 and another PT 3dA representative (BL107) showed a delay in initiation of CA4 from LBL to LGL, but not in the reverse condition, as previously reported for BL107 by these authors and characterized as ‘phenotypic group 3’. Furthermore, some PT 3dA strains, common to both studies (RS9916 and WH8020), exhibited different behaviors since these strains reached a significantly lower Exc_495:545_ at HBL than LBL in the previous but not the present study, strengthening the idea that PT 3dA cells have a more variable acclimation phenotype in HBL. In contrast, the majority of PT representatives had typical dynamics, defined by (Humily et al., 2013) as ‘phenotypic group 1’. This difference might be attributed to variability in the genomic localization of the CA4-A island, which can be found virtually anywhere in the genome for PT 3dA strains, including the 5’-end of the PBS region, while the CA4-B island is systematically located in the middle of the PBS rod region on their PT 3dB counterparts (Humily et al., 2013; Grébert et al., 2022). Certainly, the most striking outcome of the shift experiments performed in the present study was the discrepancy in acclimation pace between the two types of chromatic acclimaters, more particularly in two out of four shifts. Indeed, in the LGL to LBL shift, the PT 3dA representatives reached maximum Exc_495:545_ ratio before PT 3dB strains, while the opposite was observed in the HBL to HGL shift. These results are somehow counterintuitive since during acclimation experiments, PT 3dA cells tended to remain in their basal state (i.e., low Exc_495:545_ ratio) until the BL:GL ratio reached 75 to 100% and conversely for PT 3dB strains for which the basal state is a high Exc_495:545_ ratio (Fig. 4). Altogether, this could indicate that CA4-capable strains require a large proportion of photons opposite to their basal state to trigger CA4 process, and that once they are abruptly shifted to the light condition opposite to their basal state (i.e. 100 % BL or GL), the CA4 kinetics is faster than for the other PT (Fig. 6).

In conclusion, our study showed that PTs 3dA and 3dB exhibit subtle differences in their phenotypes, which may explain that in a spectral niche with a mix of blue and green wavelengths, they can coexist not only with blue and green light specialists, but also with their chromatic acclimater counterparts (Huisman et al., 2002; Stomp et al., 2004; Luimstra et al., 2020). Future studies should confirm these hypotheses, either using co-culture experiments of different PTs or by studying the variations of the relative abundance of PTs triggered by changes of the underwater light field.

## DATA AVAILABILITY STATEMENT

The authors confirm that the data supporting the findings of this study are available within the article and/or its supplementary materials.

## CONFLICT OF INTEREST

The authors declare no conflict of interest.

## AUTHOR CONTRIBUTIONS

LD, FP and LG conceived the experiments, analyzed the data and wrote the paper. LD, BG, JC and MR collected the samples and performed the physiological measurements. LD, BG and JC ran the flow cytometry analyses. LD made the figures. All the authors read and approved the final manuscript.

## FUNDING

This work was supported by the French “Agence Nationale de la Recherche” Programs EFFICACY (ANR-19-CE02-0019).

## ACKNOWLEDGEMENTS

We would like to thank Priscillia Gourvil, Martin Gachenot and Michele Grego from the Roscoff Culture Collection (http://roscoff-culture-collection.org/) for maintaining the *Synechococcus* strains used in this study.

## SUPPLEMENTARY MATERIAL

**Supplementary Figure 1: Representative fluorescence excitation spectra of *Synechococcus* strains RSS9916 (PT 3dA) and RS9915 (PT3 dB) grown in the various mixtures of blue and green light used in this study in low light and at 25°C. (A)** 100% BL. **(B)** 75% BL – 25% GL. **(C)** 50% BL – 50% GL. **(D)** 25% BL – 75% GL. **(E)** 100% GL. The LEDs spectra are shown in black dashed line, while the excitation spectra of RS9916 and RS9915 are represented by solid dark and light grey lines, respectively.

**Supplemental Figure 2: Average Em_650:680_ fluorescence emission ratio (a proxy of the PC:TA ratio) of six *Synechococcus* strains grown in different conditions of temperature, light quantity and quality. (A)** 18°C low light, **(B)** 18°C high light, **(C)** 25°C low light, **(D)** 25°C high light. Histograms represent average and standard deviations of triplicate measurements. Stars above histograms indicate significantly different values between PT 3dA and 3dB representatives in a given condition of light quality and temperature (Student test or Wilcoxon-Mann-Whitney test, depending on normality and variance homogeneity, p-value < 0.05).

**Supplemental Figure 3: Average flow cytometric forward scatter (a proxy of cell diameter) of six *Synechococcus* strains grown in different conditions of temperature, light quantity and quality. (A)** 18°C low light, **(B)** 18°C high light, **(C)** 25°C low light, **(D)** 25°C high light. Histograms represent average and standard deviations of triplicate measurements. Stars above histograms indicate significantly different values between PT 3dA and 3dB representatives in a given condition of light quality and temperature (Student test or Wilcoxon-Mann-Whitney test, depending on normality and variance homogeneity, p-value < 0.05).

**Supplemental Figure 4: Exc_495:550_ fluorescence excitation ratio (a proxy of PUB:PEB ratio) of the ten *Synechococcus* strains after an abrupt shift of light quality. (A, E)** LGL to LBL, **(B, F)** HGL to HBL, **(C, H)** LBL to LGL, **(D, H)** HBL to HGL. **(A, B, C, D)** PT 3dA, **(E, F, G, H)** PT 3dB. Each point represents one measurement.

## Notes

### Competing Interest Statement

The authors have declared no competing interest.

