## Supplementary figures and images for "Differential acclimation kinetics of the two forms of Type IV chromatic acclimaters occurring in marine *Synechococcus* cyanobacteria"

### Supplemental Fig. 1

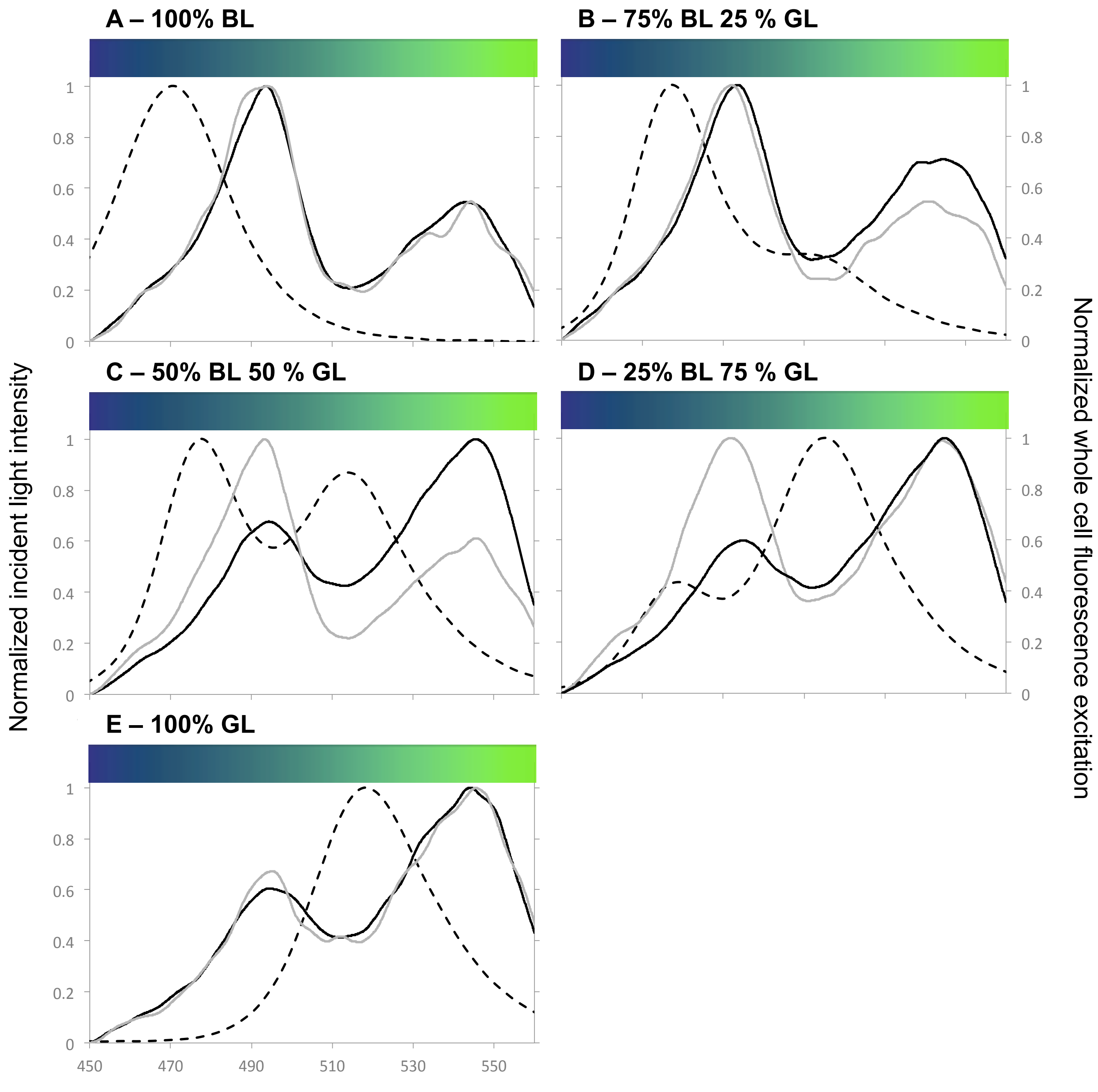

### Supplemental Fig. 2

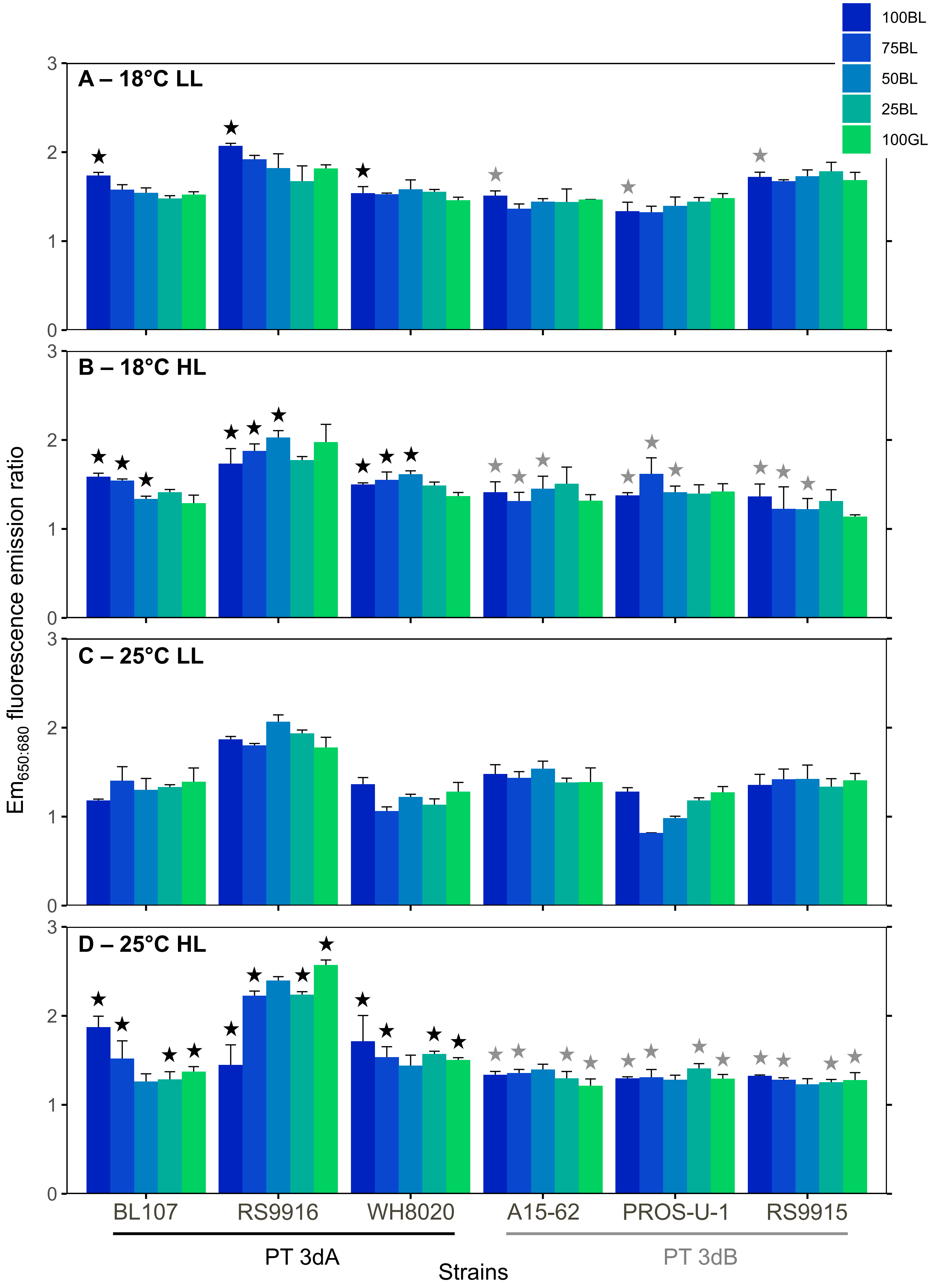

### Supplemental Fig. 3

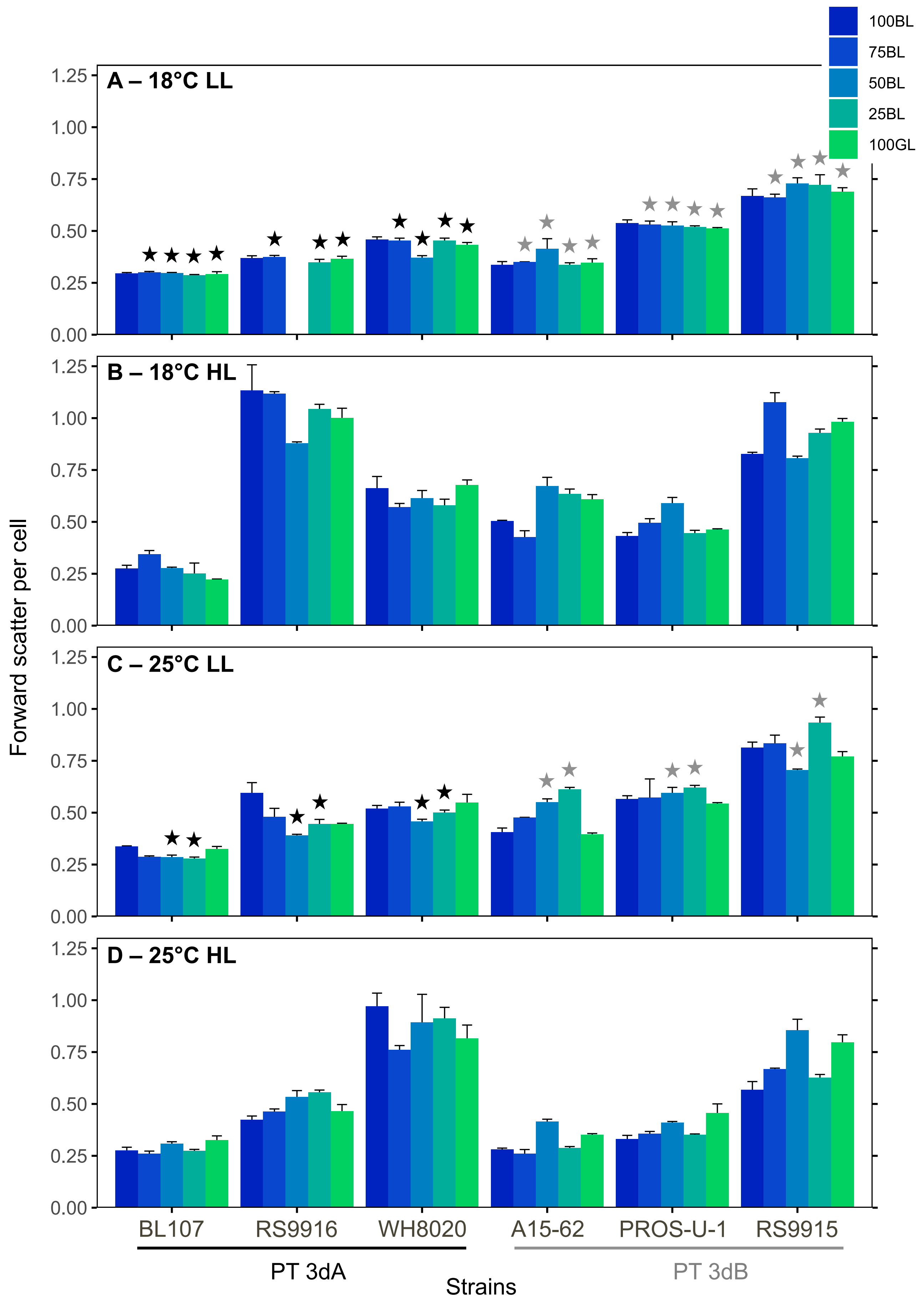

### Supplemental Fig. 4

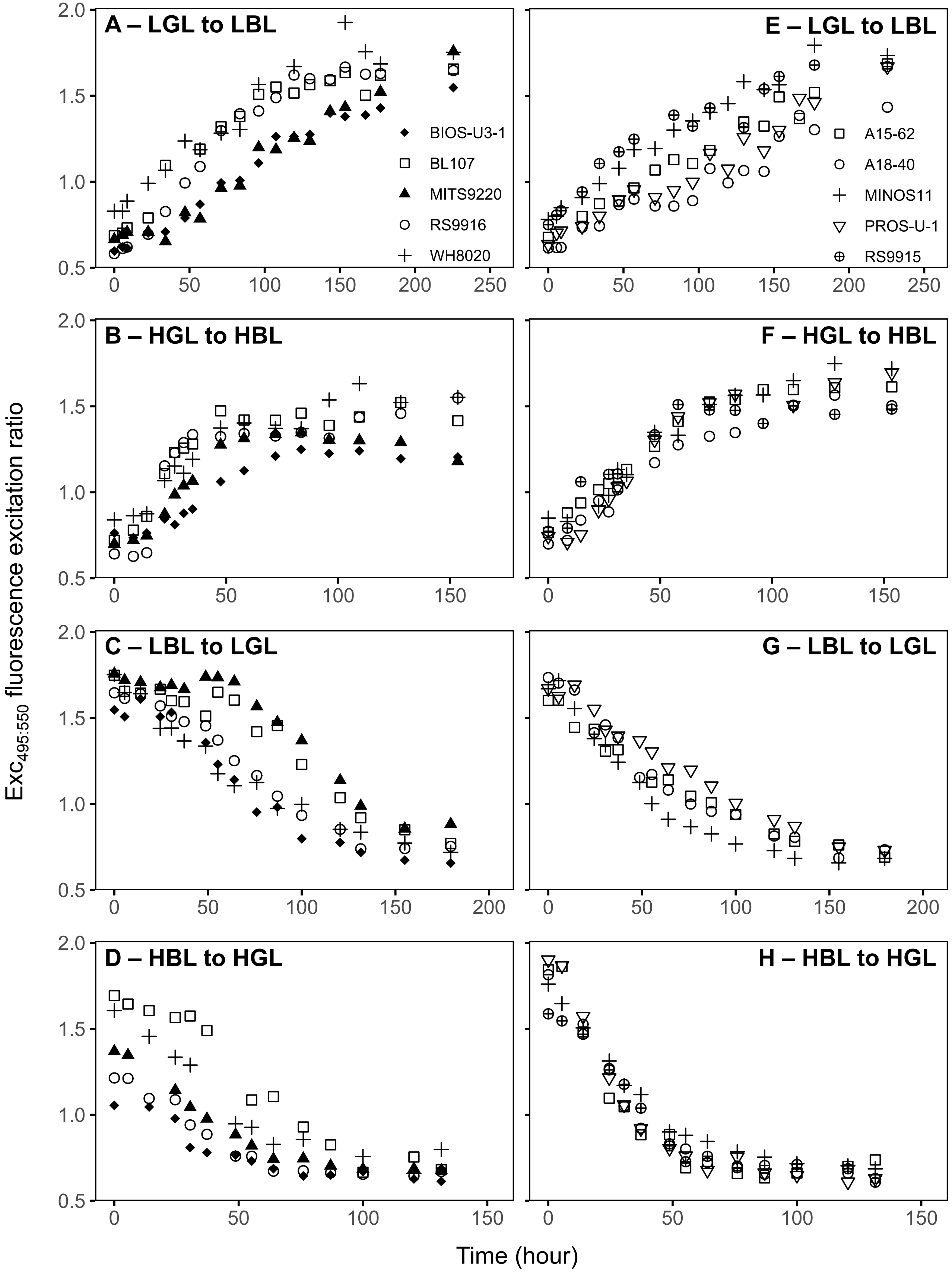
